# Brilacidin, a host defense peptide mimetic, potentiates ibrexafungerp antifungal activity against the human pathogenic fungus *Aspergillus fumigatus*

**DOI:** 10.1101/2024.04.05.588305

**Authors:** Thaila Fernanda dos Reis, Camila Diehl, Camila Figueiredo Pinzan, Patrícia Alves de Castro, Gustavo H. Goldman

## Abstract

*Aspergillus fumigatus* is the primary etiological agent of aspergillosis. Here, we show that the host defense peptide mimetic, brilacidin (BRI) can potentiate ibrexafungerp (IBX) against clinical isolates of *A. fumigatus*. CAS-resistant strains with mutations in *fks1* that encodes the 1,3-β-D-glucan synthase are not IBX-resistant and BRI+IBX can inhibit their growth. The combination of BRI+IBX plays a fungicidal role, increases the fungal cell permeability and decreases the fungal survival in the presence of A549 epithelial cells.

## Introduction

*Aspergillus fumigatus* is a saprophyte thermotolerant fungus that can cause a series of distinct clinical entities named aspergillosis, which the most severe form is the invasive pulmonary aspergillosis (IPA)^1^. Currently, recommended treatment against IPA is based on first-line azole drugs and second-line amphotericin and echinocandins, such as caspofungin^1^. Due to the increased emergence of *A. fumigatus* azole-resistant mutants^2^, new alternatives are necessary to treat aspergillosis. A recent addition to the antifungal repertoire is the triterpenoid ibrexafungerp (IBX), a semi-synthetic derivative of enfumafungin^3,4^. IBX binds to the 1,3-β-D-glucan synthase, the same target of echinocandins, but in contrast to this class of compounds, it has oral bioavailability^3,4^. Currently, IBX is approved as an oral drug for the treatment of vulvovaginal candidiasis, and it is in the process of approval against invasive candidiasis and aspergillosis^3-5^. Here, we describe brilacidin (BRI), a host defense peptide mimetic, as a potentiating agent of IBX against *A. fumigatus*.

BRI is a new chemical entity, a small molecule, constructed aiming to mimic the amphiphilic structure of host defense proteins (HDPs), having one surface with positively charged groups (cationic) and the opposite surface consisting of hydrophobic groups^6-9^. With this general synthetic form, there is no need for an agent to be of the size or composition of naturally occurring proteins to effectively function as an HDP, as the ability to act as an HDP is retained by the much smaller synthetic amphiphilic molecule. The minimal inhibitory concentration (MIC) of BRI for *A. fumigatus* was measured as higher than 80 µM and the combination of 20 µM BRI with 0.2 or 0.5 µg/mL of CAS or 0.125 and 0.25 µg/mL of VOR for 48 h at 37 °C reduced and completely inhibited conidial viability between 85 and 100 %, respectively, while these concentrations of CAS and VOR alone allowed slow conidial germination^10^. To test if BRI could potentiate the effects of another *fks1* (that encodes the 1,3-β-D-glucan synthase) inhibitor, we checked the impact of the combination between BRI and IBX against *A. fumigatus*. The viability of *A. fumigatus* was assessed by defining the number of colony forming units (CFU) after exposure of the fungal conidia to different combinations of BRI (20 and 40 µM)+IBX (0.125, 0.25, and 0.5 µg/mL) followed by plating on minimal medium. The different combinations of BRI+IBX reduced conidial viability from 98 to 100 % (**Figure 1a**). We also observed that a CAS-resistant strain DPL1035 (with a S679P *fks1* mutation) was 100 % resistant to combinations of BRI (20 and 40 µM)+IBX (0.125 µg/mL) but showed 96 to 100 % reduction in conidial viability in the presence of BRI (20 and 40 µM)+IBX (0.25 and 0.5 µg/mL) (**Figure 1a**). The Fractional Inhibitory Concentration (FIC) index for BRI+IBX was 0.82 and 0.99 indicating an additive effect against *A. fumigatus* A1163 and DPL1035 CAS-resistant isolate, respectively (**Figure 1b**).

**Figure 1.**
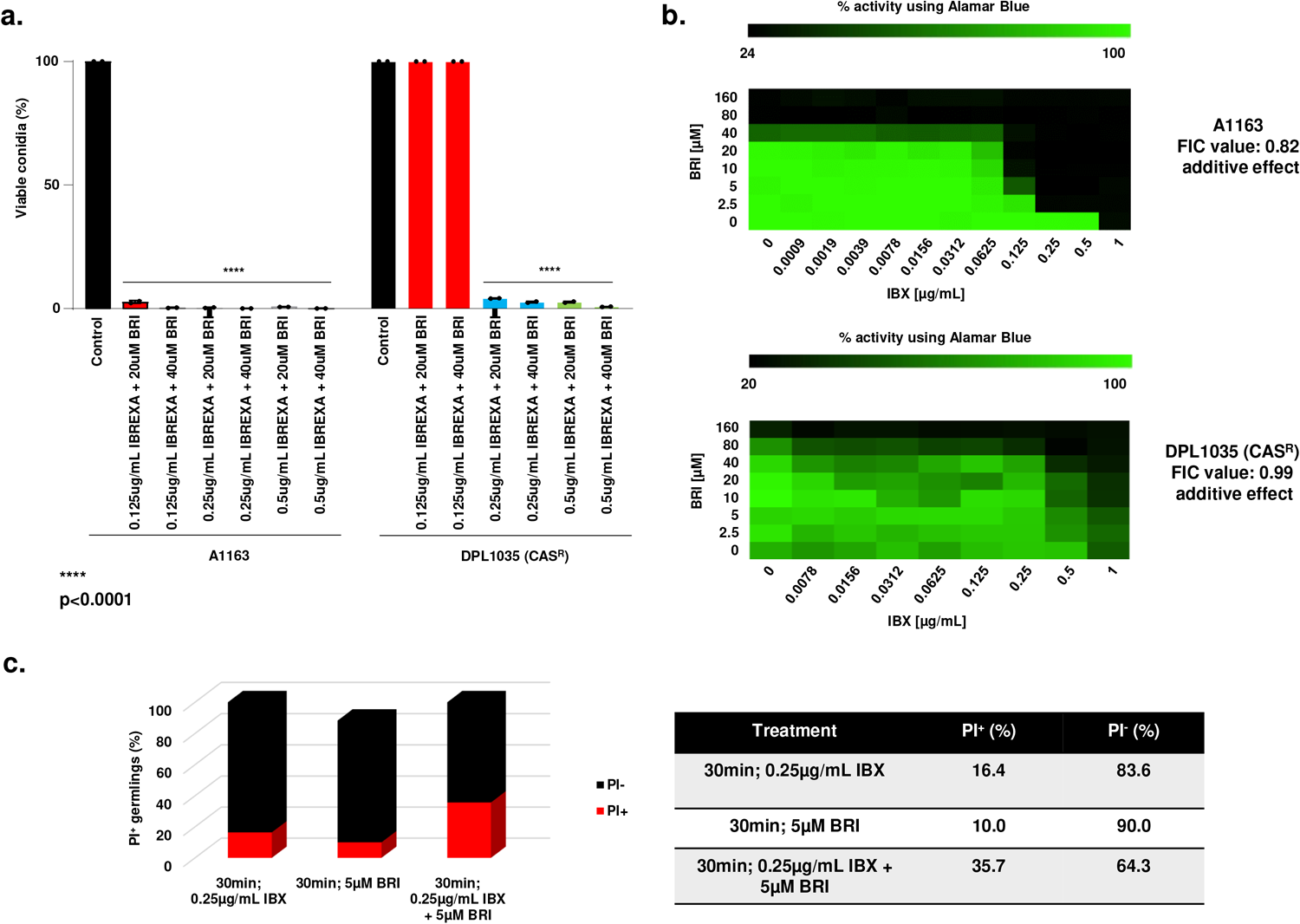
BRI+IBX has synergism against *A. fumigatus*. **a**. *A. fumigatus* conidia were incubated for 48h at 37°C with different combinations of BRI+IBX. After this period, non-germinated conidia were plated on MM, and colony-forming units (CFUs) were assessed. The results are expressed as the % of viable conidia concerning initial inoculum (n=100) and are the average of three repetitions ± SD (t-test; *p* < 0.0001). **b**. The Fractional Inhibitory Concentration (FIC) index for BRI+IBX. Results show the average of three independent experiments. **c**. BRI+IBX increases the membrane permeabilization. *A. fumigatus* was grown 16h at 30°C in MM and exposed to MM supplemented with IBX [0.25µg/mL], BRI [5µM], or CAS+BRI for 30 min. Propidium iodide (PI) was added at a 50 mg/ml concentration for 5 min. The results are expressed as the % of PI^+^ and PI^-^and are the average of three independent repetitions of 30 germlings each ± SD (n=50; t-test; *p* < 0.0001).

Antimicrobial peptides target directly or indirectly the microorganism plasma membrane disrupting their membrane potential^11,12^, and BRI has been shown to act by a similar mechanism in various (non-fungal) microorganisms^13-15^. We determined the effect of BRI+IBX on cell permeability by using propidium iodide (PI). This fluorescent DNA-binding dye freely penetrates the cell membranes of dead or dying cells but is excluded from viable cells. When *A. fumigatus* germlings were incubated with IBX for 30 min, 16.4 % were stained by PI, while exposure to BRI 5 µM yielded 10 % PI^+^ (**Figure 1c**). The combination of BRI+IBX yielded 35.7 % of PI^+^ germlings (**Figure 1c**). These results suggest that, together, BRI+IBX increases the *A. fumigatus* cell membrane permeabilization.

To get more insights about the efficacy of the BRI + IBX combination, we also evaluated if the combination of BRI+IBX could inhibit CAS-resistant and VOR-resistant *A. fumigatus* clinical isolates (**Table 1**). All these strains, including the reference strain A1163, 9 *A. fumigatus* clinical isolates VOR-resistant but susceptible to CAS (MEC CAS of 0.25 µg/mL) and two CAS-resistant clinical strains [MEC CAS of 16 µg/mL; strains DPL1035, with a known *fks1* mutation (this strain holds a S679P mutation); and strain CM7555 with unknown mutation(s)] have a MEC of 1.0 µg/mL for IBX and a MIC > 80 µM for BRI (**Table 1**). The addition of BRI at 10 µM combined with 0.25 µg/mL of IBX completely inhibited the growth of all tested strains, including those that are resistant to CAS or with known resistance to azoles (with the TR34/L98H mutation; **Table 1**). These data suggest that BRI potentiates IBX activity against CAS- or VOR-resistant strains of *A. fumigatus*.

**Table 1.** MECs e MICs of *A. fumigatus* strains.

| Strains | MEC CAS (µg/mL) | MEC IBX (µg/mL) | MIC BRI (µM) | BRI (10 µM)+ IBX (0.25 µg/mL)* | Origin |
| --- | --- | --- | --- | --- | --- |
| A1163 | 0.25 | 1.0 | > 80 | - | FGSC** |
| DPL1035 | 16.0 | 1.0 | > 80 | - | David Perlin lab |
| CM7555 | 16.0 | 1.0 | > 80 | - | Gustavo Goldman lab |
| CYR15-184 | 0.25 | 1.0 | > 80 | - | Katrien Lagrou lab |
| CYR15-190 | 0.25 | 1.0 | > 80 | - | Katrien Lagrou lab |
| CYR15-192 | 0.25 | 1.0 | > 80 | - | Katrien Lagrou lab |
| CYR15-195 | 0.25 | 1.0 | > 80 | - | Katrien Lagrou lab |
| CYR15-202 | 0.25 | 1.0 | > 80 | - | Katrien Lagrou lab |
| CYR15-212 | 0.25 | 1.0 | > 80 | - | Katrien Lagrou lab |
| CYR15-213 | 0.25 | 1.0 | > 80 | - | Katrien Lagrou lab |
| CYR15-215 | 0.25 | 1.0 | > 80 | - | Katrien Lagrou lab |
| CYR15-220 | 0.25 | 1.0 | > 80 | - | Katrien Lagrou lab |
| CYR15-221 | 0.25 | 1.0 | > 80 | - | Katrien Lagrou lab |
| CYR15-222 | 0.25 | 1.0 | > 80 | - | Katrien Lagrou lab |
| CYR15-224 | 0.25 | 1.0 | > 80 | - | Katrien Lagrou lab |
| CYR15-225 | 0.25 | 1.0 | > 80 | - | Katrien Lagrou lab |
| CYR15-226 | 0.25 | 1.0 | > 80 | - | Katrien Lagrou lab |
| CYR15-228 | 0.25 | 1.0 | > 80 | - | Katrien Lagrou lab |
| CYR15-229 | 0.25 | 1.0 | > 80 | - | Katrien Lagrou lab |
| CYR15-230 | 0.25 | 1.0 | > 80 | - | Katrien Lagrou lab |
| CYR15-231 | 0.25 | 1.0 | > 80 | - | Katrien Lagrou lab |
| CYR15-109 | 0.25 | 1.0 | > 80 | - | Katrien Lagrou lab |
| CYR15-147 | 0.25 | 1.0 | > 80 | - | Katrien Lagrou lab |

\*, -, no growth; \*\*,FGSC, Fungal Genetics Stock Center (www.fgsc.net)

Further we used the A549 pulmonary cells to check the *in vitro* toxicity played by BRI+IBX. Initially, the A549 cells were incubated with 40 µM of BRI with or without increasing IBX concentrations for 24 h, after which cell viability was assessed by XTT assay (**Figure 2a**). As a positive control, we used DMSO 10%, which reduced cell viability by 90%. Neither BRI, IBX, nor their combinations significantly reduced cell viability compared to the control (**Figure 2a**). To determine the killing rates, A549 cells were challenged with 1:10 ratios (A549-conidia), and we observed a significant reduction of more than 25, 25, and 65 % in the fungal viability in BRI 40 µM, IBX 0.25 µg/mL, and BRI (40 µM)+IBX (0.25 µg/mL), respectively (**Figure 2b**). VOR, the standard of care in controlling *A. fumigatus* cell growth, showed a comparable growth reduction to the combination of BRI+IBX (**Figure 2b**).

**Figure 2.**
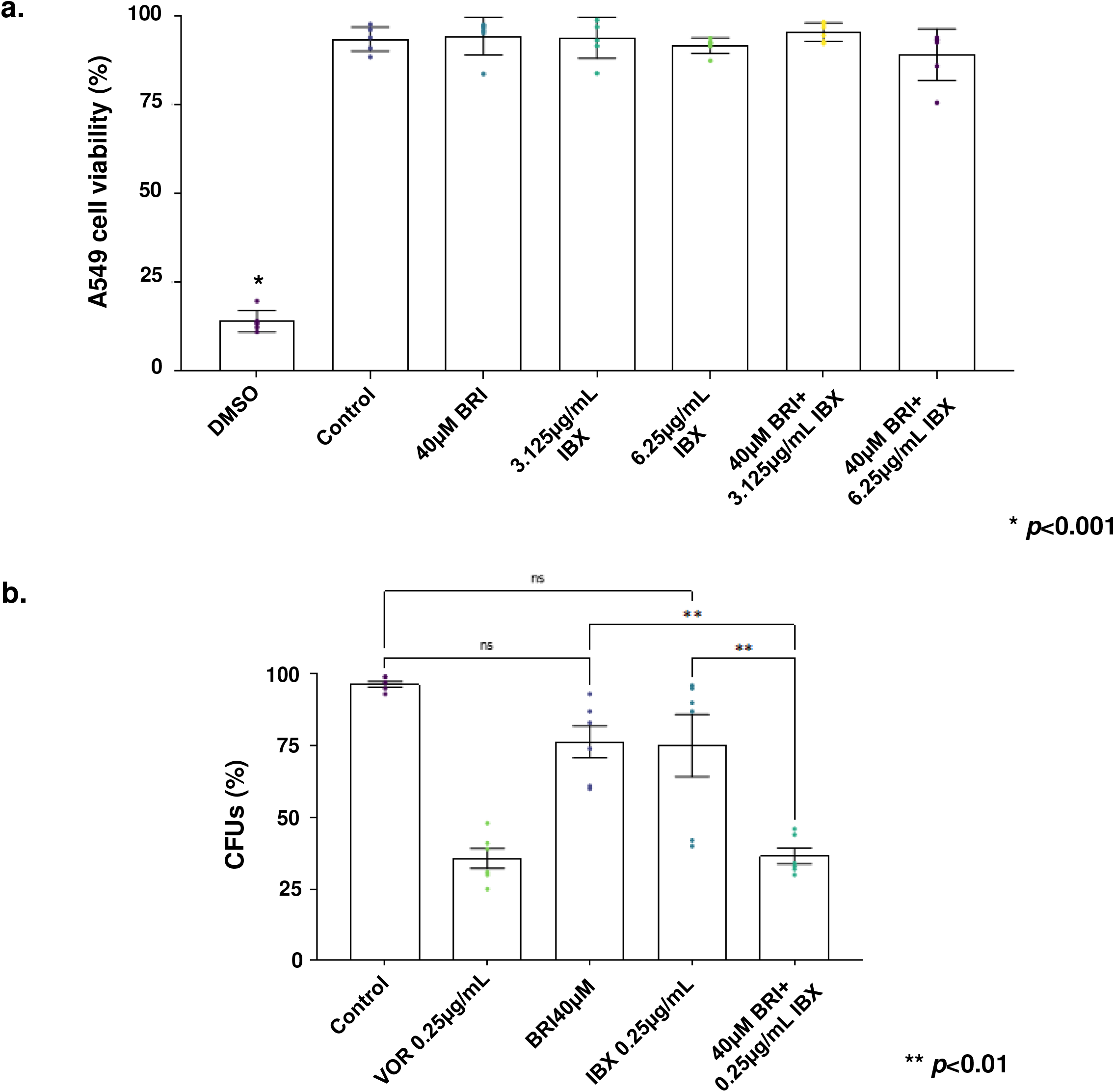
The combination of BRI+IBX is not toxic and reduces the *A. fumigatus* fungal burden during the infection of A549 lung epithelial cells. **a**. The cytotoxicity of BRI, IBX or BRI + IBX was assessed by exposure of A549 lung epithelial cells to BRI [40µM], IBX [3.125µg/mL or 6.25µg/mL], BRI [40µM] + IBX [3.125µg/mL or 6.25µg/mL] for 24h followed by the measurement of the cell metabolic (as determined by XTT). DMSO 10% and fresh conidia were used as controls. The percentage of cell viability is expressed as the absorbance value of the experiment well/absorbance value of the control well x 100. **b**. A549 lung epithelial cells were seeded at a density of 10^6^ cells/mL and challenged with *A. fumigatus* conidia at a multiplicity of infection (MOI) of 1:10 in the absence or presence of BRI [40µM], IBX [0.25µg/mL] or BRI [40µM] + IBX [0.25µg/mL]. After 24 h of incubation, the media was removed, and the cell suspensions were plated on solid Sabouraud Dextrose media. The number of CFUs was determined after 24 hours of growth. The CFU percentage for each sample was calculated compared to untreated cells (Control), and the results were plotted using GraphPad Prism (GraphPad Software, Inc., La Jolla, CA, USA). The fungicidal drug VOR was included as a control. All the results are the average of three repetitions ±SD. \*\**p*-value<0.01.

Interestingly, one of the *A fumigatus* CAS-resistant mutants that have a mutation in the *fks1* gene is not resistant to IBX (**Table 1**), suggesting that this mutation at S679P is not able to confer IBX-resistance. These results are similar to *Candida* spp. because IBX’s binding site seems to be partially divergent from that of the echinocandins since IBX is active against echinocandin-resistant *Candida* isolates^16,17^.

Taken together, these results indicate that the combination BRI+IBX is fungicidal, can increase the membrane permeability, has increased fungicidal activity in the presence of A549 lung epithelial cells, and wholly overcomes CAS-resistance in echinocandin-resistant *A. fumigatus* clinical isolates and VOR-resistant isolates. In summary, our work with BRI+IBX brings an essential contribution to the combination therapies already proposed for IBX, such as IBX+voriconazole, IBX+isavuconazole, and IBX+amphotericin that were synergistic against both *A. fumigatus* azole-susceptible and –resistant strains^18-20^. We extended the previously observed number of antifungal drugs BRI can potentiate against *A. fumigatus*.

## Acknowledgments

We thank the Fundação de Amparo à Pesquisa do Estado de São Paulo (FAPESP) grant numbers 2022/08796-8 (CD), 2021/04977-5 (GHG), and 2021/10599-3 (The Antimicrobial Resistance Institute of São Paulo, The Aries Project), the Conselho Nacional de Desenvolvimento Científico e Tecnológico (CNPq), FAPESP and Fundação Coordenação de Aperfeiçoamento do Pessoal do Ensino Superior (CAPES) grant number 405934/2022-0 (The National Institute of Science and Technology INCT Funvir), and CNPq 301058/2019-9 from Brazil to GHG, and the National Institutes of Health/National Institute of Allergy and Infectious Diseases (R01AI153356) from the USA (G.H.G.). This work was funded by the Joint Canada-Israel Health Research Program, jointly supported by the Azrieli Foundation, Canada’s International Development Research Centre, Canadian Institutes of Health Research, and the Israel Science Foundation (GHG). We also thank David Perlin and Katrien Lagrou for sending *A. fumigatus* caspofungin- and voriconazole-resistant strains, and Innovation Pharmaceuticals Incorporation, Scynexis and GlaxoSmithKline for providing brilacidin and ibrexafungerp, respectively.

